# The pd-l1/pd-1 axis blocks neutrophil cytotoxicity in cancer

**DOI:** 10.1101/2020.02.28.969410

**Authors:** Maya Gershkovitz, Olga Yajuk, Tanya Fainsod-Levi, Zvi Granot

## Abstract

The PD-L1/PD-1 axis was shown to promote tumor growth and progression via the inhibition of anti-tumor immunity. Blocking this axis was shown to be beneficial in maintaining the anti-tumor functions of the adaptive immune system. Still, the consequences of blocking the PD-L1/PD-1 axis on innate immune responses remain largely unexplored. In this context, neutrophils were shown to consist of different subpopulations, which possess either pro- and anti-tumor properties. PD-L1-expressing neutrophils are considered pro-tumor as they can suppress cytotoxic T-cells. That said, we found that PD-L1 expression is not limited to tumor promoting neutrophils but is also evident in anti-tumor neutrophils. We show that neutrophil cytotoxicity is effectively and efficiently blocked by tumor cell expressed PD-1. Furthermore blocking of either neutrophil PD-L1 or tumor cell PD-1 maintains neutrophil cytotoxicity resulting in enhanced tumor cell apoptosis. Importantly, we show that tumor cell PD-1 blocks neutrophil cytotoxicity and promotes tumor growth via a mechanism independent of adaptive immunity. Taken together, these findings highlight the therapeutic potential of enhancing anti-tumor innate immune responses via blocking of the PD-L1/PD-1 axis.

## INTRODUCTION

Anti-tumor immune surveillance is an important feature of the immune system where immune cells recognize and eliminate malignancies (Chen and Mellman, 2013; Hanahan and Coussens, 2012; Nanda and Sercarz, 1995). However, immune cells make a significant proportion of the tumor microenvironment (TME) and they were shown to contribute to tumor development and progression (Barberis et al., 2016). This dramatic change in immune function is mediated via various evasion mechanisms (Drake et al., 2006; Ribas, 2015), one of which is the expression of immune checkpoint molecules by tumor cells. Under physiological conditions, the expression of immune checkpoint molecules regulates immune responses and prevents autoimmunity (Sharma and Allison, 2015; Topalian et al., 2015). In cancer, the expression of these molecules enables blocking of anti-tumor immunity and promotes immune evasion (Ribas, 2015).

The programmed cell death protein 1 (PD-1; also known as Pdcd1) receptor and its ligand PD-1 ligand 1 (PD-L1) serve as an immune checkpoint pathway and are of significant clinical importance (Chikuma, 2016; Trivedi et al., 2015). Tumor cells benefit from the expression of PD-L1 as its expression inhibits the proliferation and activation of cytotoxic T cells (Amarnath et al., 2011; Blank et al., 2004; Butte et al., 2007). Interestingly, the expression of the inhibitory ligand, PD-L1, is not limited only to tumor cells in the TME (Gibbons Johnson and Dong, 2017). Neutrophils in the TME and in the peritumoral tissue also express high levels of PD-L1 (He et al., 2015; Wang et al., 2017). Tumor-infiltrating neutrophils were shown to play important, yet multifaceted and sometimes opposing roles (Sionov et al., 2015b). The tumor promoting properties of neutrophils were widely described as they have the ability to initiate tumorigenesis (Katoh et al., 2013), enhance tumor progression via direct stimulation of tumor cells proliferation (Houghton et al., 2010), enhance angiogenesis (Jablonska et al., 2010; Nozawa et al., 2006), suppress anti-tumor immune effector cells (de Kleijn et al., 2013; Gabrilovich et al., 2012) and promote metastatic spread (Casbon et al., 2015; Cools-Lartigue et al., 2013; Spicer et al., 2012). In contrast, studies have shown that neutrophils may also possess anti-tumor features as they can kill tumor cells via the secretion of H_2_O_2_ (Gershkovitz et al., 2018a; Granot et al., 2011). The role of PD-L1^pos^ neutrophils is associated with a tumor-promoting phenotype, they were found to suppress cytotoxic T cells, thereby enhance disease progression and limit patient survival (de Kleijn et al., 2013; He et al., 2015).

Although the expression of PD-L1 is mainly attributed to tumor cells and PD-1 expression is attributed to T cells, the expression pattern of PD-L1/PD-1 in cancer is more complex (Gibbons Johnson and Dong, 2017; Simon and Labarriere, 2017). A study by Kleffel and colleges (Kleffel et al., 2015) showed that murine and human melanomas contain PD-1 expressing cancer subpopulations and demonstrate that melanoma PD-1 cell-intrinsic signaling promotes tumorigenesis. Although the role of PD-L1 on neutrophils is not fully understood, this observation raises the possibility that PD-L1^pos^ neutrophils can directly interact with PD-1^pos^ tumor cells. In this study, we examined the role played by the PD-L1/PD-1 axis in regulating neutrophil cytotoxicity toward tumor cells. We found that neutrophil PD-L1 not only acts to suppress T cell responses, but can also directly inhibit neutrophil cytotoxicity. Blocking the PD-L1/PD-1 interaction enhances tumor cells susceptibility to neutrophil cytotoxicity. In a previous study (Granot et al., 2011), we showed that neutrophils possess an anti-metastatic role when encountering tumor cells in the pre-metastatic niche. In line with these observations, we found that PD-1 levels in tumor cells have a significant impact on the anti-metastatic function of neutrophils. While neutrophils efficiently eliminate control tumor cells, PD-1 over-expressing tumor cells are resistant to neutrophil cytotoxicity. Our observations provide a wider scope to the role PD-L1 plays on neutrophils in cancer, and highlight the therapeutic potential of blocking the PD-L1/PD-1 axis not only in propagating anti tumor T cell responses but also in facilitating neutrophil cytotoxicity.

## RESULTS

### Immature low-density neutrophils express higher levels of PD-L1

The PD-L1/PD-1 immune checkpoint is well characterized in the context of cancer where PD-L1-expressing tumor cells were shown to functionally inhibit the cytotoxicity of PD-1-expressing T cells. Furthermore, blocking of PD-L1/PD-1 interaction results in the maintenance of T cells anti-tumor cytotoxicity and the consequent reduction in tumor mass. Although the expression of PD-L1 is mainly attributed to tumor cells, PD-L1 is also expressed by various immune cells including lymphocytes, monocytes and granulocytes (**Fig. 1A**).

**Figure 1.**
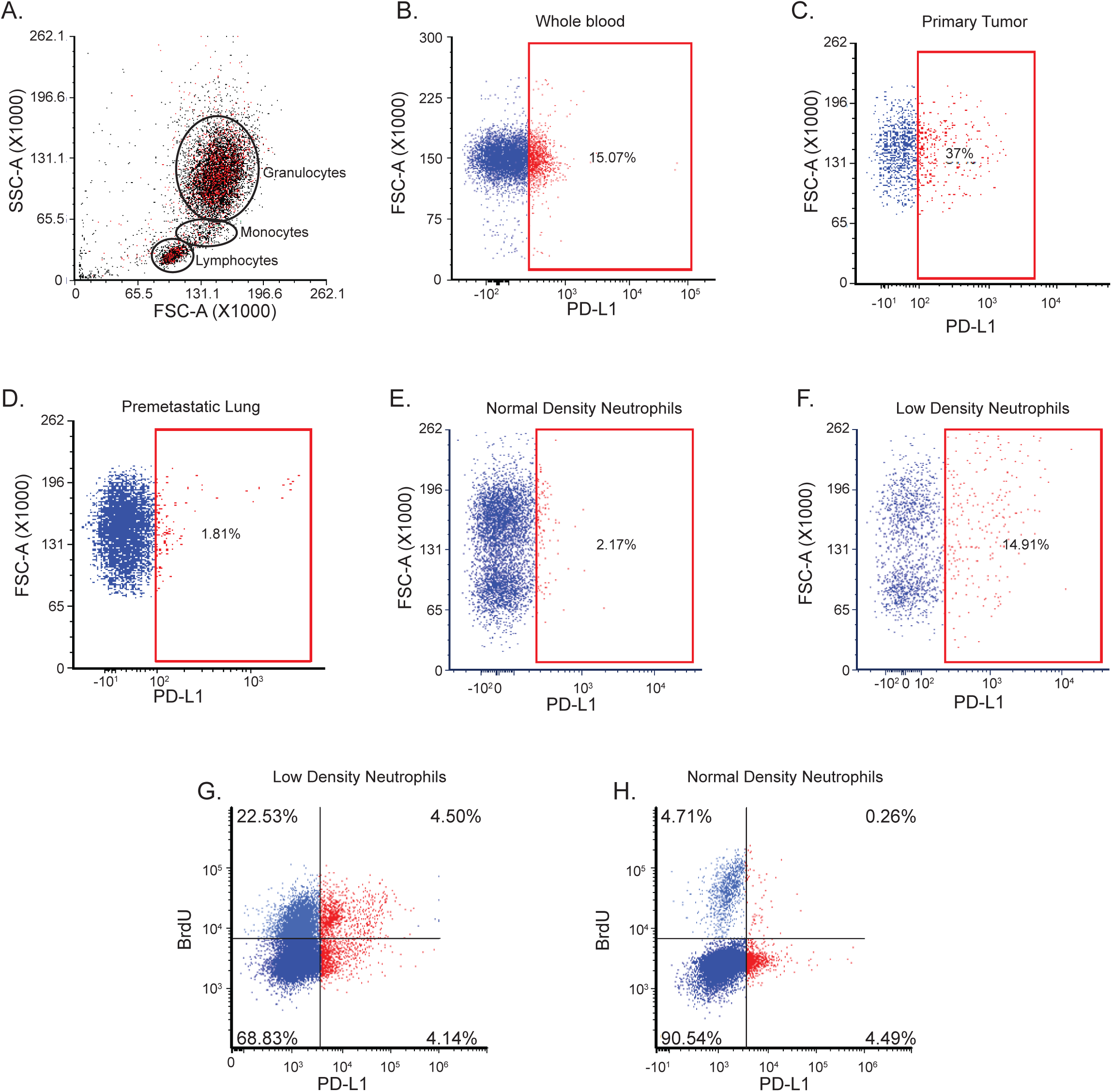
PD-L1^pos^ neutrophils are associated with immature, low-density phenotype. **A**. FACS analysis of PD-L1^pos^ cells (red) in whole blood from a tumor-bearing mouse. **B-D**. FACS analysis of PD-L1 expression in Ly6G^+^ neutrophils isolated from the circulation (B), the primary tumor (C) and the premetastatic lung (D) of a tumor-bearing mouse. Red gate represents the PD-L1^pos^ subpopulation. **E-F**. FACS analysis of PD-L1 expression in Ly6G^+^ neutrophils normal-density neutrophils (E) and low-density neutrophils (F) from a tumor-bearing mouse, red gate represents PD-L1^pos^ subpopulation. **G-H**. FACS analysis of BrdU staining in circulating Ly6G^+^ low-density neutrophils (G) and normal-density neutrophils (H) from a 4T1 tumor-bearing mouse. Red gate represents PD-L1^pos^ subpopulation.

We previously showed that the neutrophil anti-tumor activity is context dependent and can be either manifested or inhibited in different tumor microenvironments (circulation, primary tumor, pre metastatic and metastatic sites). Specifically, we showed that tumor associated neutrophils (TAN) do not exhibit anti-tumor functions whereas circulating neutrophils or neutrophils associated with the premetastatic niche exhibit high anti-tumor cytotoxicity (Gershkovitz et al., 2018b). To gain insight into the role PD-L1 plays in regulating neutrophil function in cancer, we evaluated the expression of PD-L1 on neutrophils isolated from the circulation or the primary tumor. On average, 15% of circulating Ly6G^+^ neutrophils and ∼40% of TAN were PD-L1^pos^ (**Fig. 1B-C**) Notably, only a negligible fraction of neutrophils (∼1.8%) isolated from the premetastatic lung were PD-L1^pos^ (**Fig. 1D**). As we have shown before, neutrophils may be divided into Normal and Low Density subsets (NDN and LDN respectively, (Sagiv et al., 2015)). As LDN were shown to possess pro-tumorigenic properties, we hypothesized that PD-L1 expression on neutrophils may be associated with this specific neutrophil subset. Indeed, while NDN consists of a small PD-L1^pos^ subset (∼2%), a significantly higher proportion of LDN are PD-L1^pos^ (∼15% compare **Fig. 1E and 1F**). LDN are heterogeneous and consist of both mature and immature neutrophils. Since immature neutrophils were shown to possess pro-tumor properties (Sagiv et al., 2015; Yang et al., 2011) we sought to determine whether PD-L1^pos^ neutrophils exhibit a more immature phenotype. To this end we treated tumor-bearing mice with a single dose of BrdU and assessed the fraction of BrdU^+^ neutrophils 24 hrs following BrdU administration (a time point where only immature cells are BrdU^+^, (Sagiv et al., 2015)). In the low-density fraction, 16.6% of the BrdU^+^ (immature) neutrophils were PD-L1^pos^ compared with only 5.6% of mature (BrdU^-^) neutrophils (**Fig. 1G**). The NDN fraction, which contained a small subpopulation of immature cells, consisted of a low percentage of PD-L1^pos^ neutrophils (∼5%), with no significant difference between the mature and the immature subpopulations (**Fig. 1H**).

To gain insight into their functional differences, we isolated PD-L1^pos^ and PD-L1^neg^ neutrophils (**Fig. 2A**). We found that oxidative burst (the production of H_2_O_2_) was significantly higher in the PD-L1^neg^ neutrophils (**Fig. 2B**). In addition, we noticed that PD-L1^pos^ cells contain a larger proportion of morphologically immature neutrophils (**Fig. 2C-D**). To test whether PD-L1^neg^ neutrophils exhibit a more pro-inflammatory phenotype compared with PD-L1^pos^ neutrophils we used the Zymosan-A induced mouse model of peritonitis. This model simulates self-resolving acute inflammation where peritoneal neutrophil numbers peak at 3 hours post Zymosan-A injection and gradually decrease over a period of 72 hours. We have previously shown that the phenotype of the peritoneal neutrophils changes from pro to anti-inflammatory towards resolution of the inflammatory response (Sagiv et al., 2015). Accordingly, we used this model to assess whether the anti-inflammatory, presumably more suppressive neutrophils, express higher levels of PD-L1. As expected, the percentage of peritoneal Ly6G^+^ neutrophils gradually decreased towards resolution (**Fig. 2E**). Notably, although the percentage of neutrophils gradually decreased, the fraction of PD-L1^pos^ neutrophils increased with time (**Fig. 2F**). These results show that the transition of neutrophil function, from pro-inflammatory to pro-resolution, is associated with an increase in the proportion of PD-L1^pos^ neutrophils and may indicate a possible immunosuppressive effect of the PD-L1^pos^ neutrophil subpopulation.

**Figure 2.**
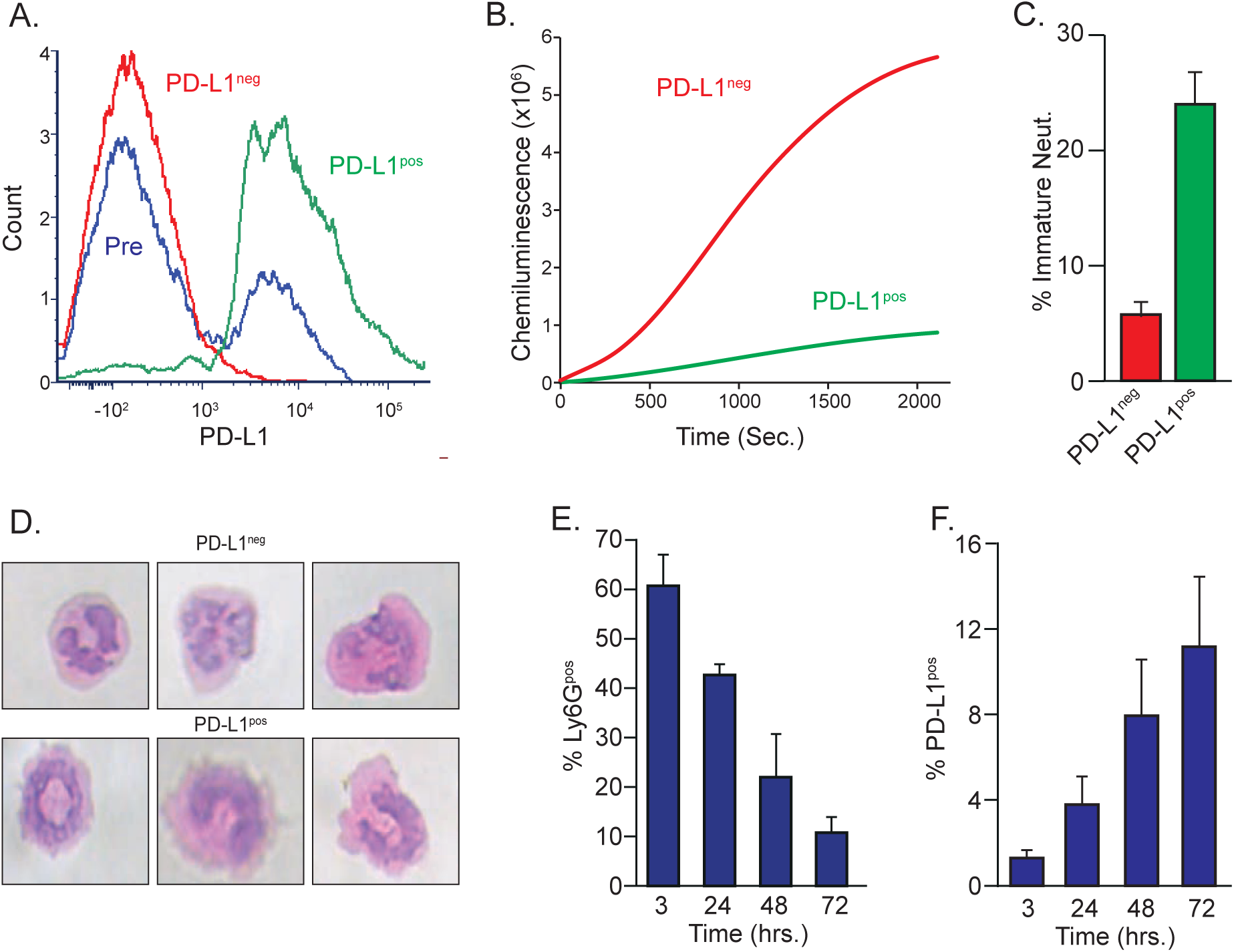
PD-L1^pos^ neutrophils are associated with the resolution phase in inflammation. **A**. FACS analysis PD-L1 expression in presorted neutrophils (Pre), PD-L1^neg^ neutrophils (red) and PD-L1^pos^ neutrophils (green). **B**. Spontaneous ROS production in PD-L1^neg^ (red) and PD-L1^pos^ (green) neutrophils. **C**. Percent of immature cells in PD-L1^neg^ (red) and PD-L1^pos^ (green) neutrophil population. **D**. Representative images of H&E staining of PD-L1^pos^ and PD-L1^neg^ neutrophils. **E**. Time dependent changes in the fraction of peritoneal neutrophils (of all CD45^+^ cells) in self-resolving peritonitis. **F**. Time dependent changes in the fraction of peritoneal of PD-L1^pos^ neutrophils (of all Ly6G^+^ cells) in self-resolving peritonitis.

### PD-1-expressing tumor cells inhibit neutrophil cytotoxicity via PD-L1/PD-1 axis

The notion that PD-L1 expression on neutrophils promotes pro-tumor features was explored and established in previous studies (Zhang and Xu, 2017). However, the effect of the PD-L1/PD-1 axis on neutrophil anti-tumor cytotoxicity was never examined. To determine the direct consequence of PD-1/PD-L1 axis on neutrophils cytotoxicity, we assessed the expression levels of PD-1 on both neutrophils and tumor cells. We found that while neutrophils do not express PD-1 (**Fig. S1**), a significant subpopulation of both 4T1 and AT3 breast cancer cell lines do express PD-1 (**Fig. 3A-B**). Importantly, we found that in a co-culture setting, specific blocking of PD-L1 enhanced the susceptibility of both 4T1 and AT3 cells to neutrophil cytotoxicity (**Fig. 3C-D**). This observation suggests that PD-1 expression on tumor cells can inhibit neutrophil cytotoxicity via PD-L1/PD-1 axis. However, this experimental platform suffers from two significant flaws: First, the PD-L1 antibody can directly affect PD-L1 expressing tumor cells. Second, the addition of an antibody to the culture may result in neutrophil killing of tumor cells via antibody-dependent cellular cytotoxicity (ADCC). To overcome these difficulties, we isolated a pure population of PD-L1^neg^ neutrophils and tested their cytotoxic capacity. Using this approach, we were able to simultaneously eliminate possible PD-L1/PD-1 interactions and avoid the presence of an antibody in the culture. We found that when co-cultured with tumor cells, PD-L1^neg^ neutrophils exhibit a significantly higher cytotoxic ability compared with the mixed population (**Fig. 3E-F**) independently corroborating the results obtained by PD-L1 blocking antibody. Finally, we used PD-1 specific shRNA to knockdown PD-1 expression in both 4T1 and AT3 cells (**Fig. 3G-H**). We then cocultured control and PD-1kd cells with cytotoxic neutrophils and found that PD-1kd cells are more susceptible to neutrophil cytotoxicity (**Fig. 3I-J**). Taken together, our observations suggest that tumor cell PD-1 limits neutrophil cytotoxicity and that PD-L1^neg^ neutrophils have enhanced cytotoxic potential.

**Figure 3.**
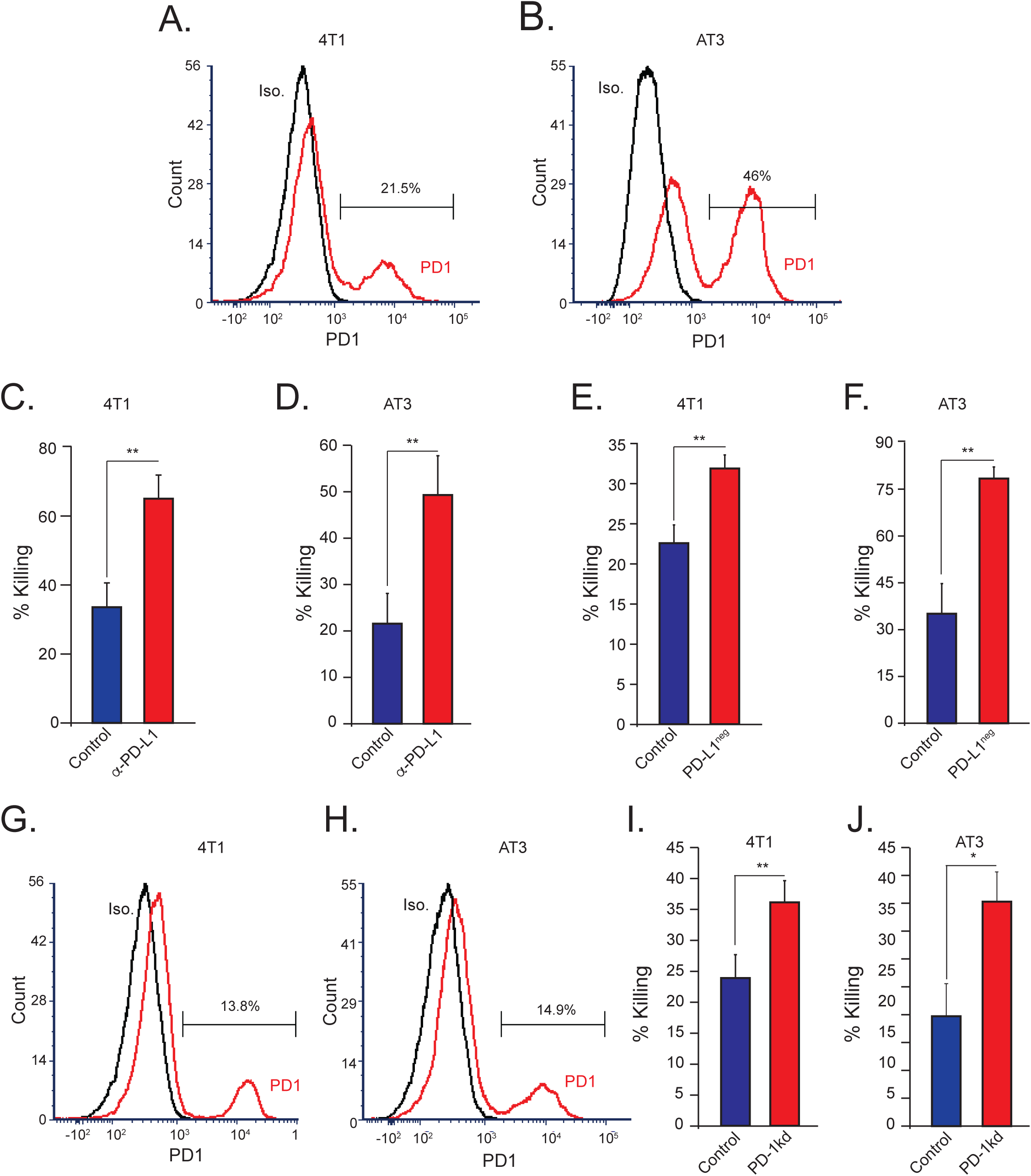
Blocking the PD-1/PD-L1 axis increases tumor cell susceptibility to neutrophil cytotoxicity. **A**. FACS analysis of PD-1 expression in 4T1 breast tumor cells. **B**. FACS analysis of PD-1 expression in AT3 tumor cells. **C-D**. Neutrophil mediated killing of 4T1 (C) and AT3 (D) tumor cells in the absence (cont.) or presence of PD-L1 blocking antibody. **E-F**. Neutrophil mediated killing of 4T1 (E) and AT3 (F) tumor cells with unsorted neutrophils (cont.) or PD-L1^neg^ neutrophils **G-H**. FACS analysis of PD1 expression in PD-1kd 4T1 (G) and AT3 (H) cells tumor (red). **I-J**. Neutrophil mediated killing of control and PD-1kd 4T1 (I) and AT3 (J) tumor cells. *p<0.05, **p<0.01.

### Tumor cell PD-1 inhibits neutrophil cytotoxicity and increases metastatic potential

To gain insight into the consequences of PD-1 expression on neutrophil function in the context of cancer we tested how tumor growth and metastatic progression are affected by overexpression of PD-1 (**Fig. 4A**). Notably, PD-1 plays a critical role in mediating the interaction between tumor cells and lymphocytes which may interfere with proper interpretation of the experiments. To overcome this difficulty we assessed tumor growth and metastatic progression in orthotopically (mammary fat pad) injected control *vs*. PD-1 overexpressing 4T1 tumor cells in NOD-SCID mice. Our data show that PD-1 overexpressing 4T1 tumors exhibited a more aggressive phenotype as they grew faster (**Fig. 4B**) and had a significantly higher metastatic load (**Fig. 4C-D**) compared to control tumors. Since these mice lack adaptive immunity, these observations highlight the inhibitory effect of the PD-L1/PD-1 axis on innate immune cells. However, since this effect on tumor growth and metastatic progression may be mediated by cells other than neutrophils, we tested the consequences of neutrophil depletion on metastatic seeding of control and PD-1 overexpressing tumor cells. First, we orthotopically injected NOD-SCID mice with control 4T1 tumor cells in order to generate a primary tumor and prime the immune system. Next, we injected control or PD-1 overexpressing 4T1^GFP+^ tumor cells via the tail vein to control or neutrophil-depleted tumor-bearing mice (**Fig. 4E**). Corroborating previous observations, neutrophil depletion (**Fig. 4F**) enhanced the seeding of control 4T1^GFP+^ tumor cells in the lungs (**Fig. 4G-H**). In contrast, neutrophil depletion had no significant effect on the seeding of PD-1 over-expressing 4T1^GFP+^ tumor cells (**Fig. 4G and 4I**). To broaden the scope of these observations, we implemented the same approach and overexpressed PD-1 in AT3 cells (**Fig. 4J**). As in 4T1 cells, neutrophil depletion increased the seeding of control tumor cells in the lungs (**Fig. 4K**) but had no effect on metastatic seeding of PD-1 overexpressing cells (**Fig. 4K**). These observations suggest that PD-1 expression on tumor cells can alter neutrophil function and inhibit their anti-tumor cytotoxicity.

**Figure 4.**
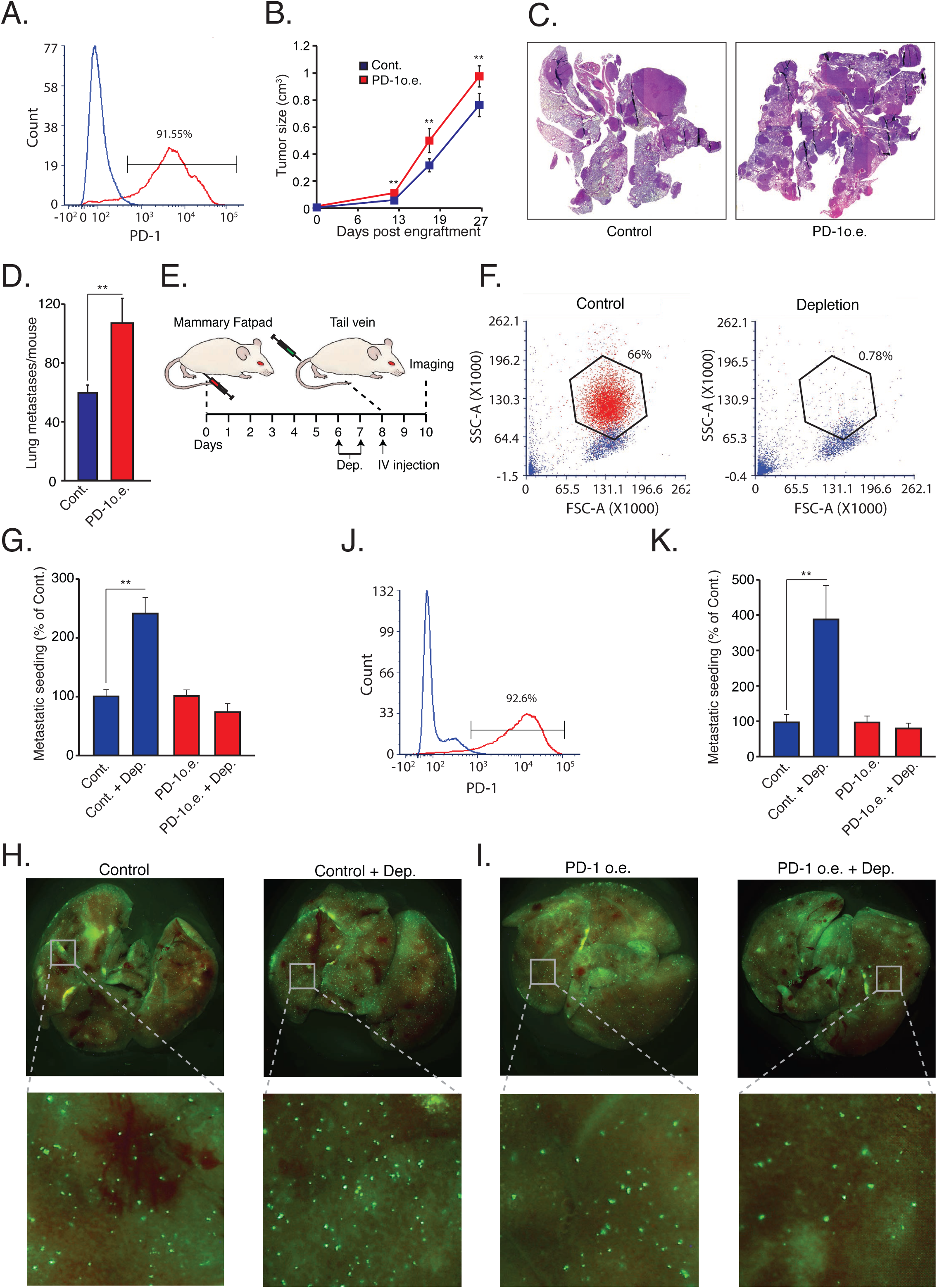
PD-1 overexpression impairs neutrophil cytotoxicity in NOD-SCID mice. **A**. FACS analysis of PD-1 overexpression in 4T1 tumor cells. **B**. Tumor growth of orthotopically injected control and PD-1 overexpressing 4T1 cells (n=5). **C**. Representative H&E staining of lungs from mice injected with control or PD-1 overexpressing 4T1 cells (n=5). **D**. Total number of spontaneous metastases in control and PD-1 overexpressing 4T1 tumor bearing mice (n=5). **E**. Experimental metastatic seeding assay - female NOD-SCID mice were orthotopically (mammary fat pad) injected with control (4T1 or AT3) tumor cells. Starting on day 6 the mice were injected daily (arrows) with either control IgG or neutrophil depleting anti-Ly6G antibody *i*.*p*. On day 8 the mice were injected with 5×10^5^ GFP^+^ control or PD-1 overexpressing cells (4T1 or AT3) via the tail vein. On day 10 the number of lung associated GFP^+^ cells were analyzed by fluorescent microscopy. **F**. FACS analysis of circulating Ly6G^+^ neutrophils in control and neutrophil depleted tumor-bearing mice. **G**. Average number of lung-associated GFP^+^ control and PD-1 overexpressing 4T1 tumor cells in control and neutrophil depleted mice (n=5). **H**. Representative images of lung associated control GFP^+^ cells in control mice (control) and neutrophil depleted mice (control + dep.). **I**. Representative images of lung associated PD-1 overexpressing GFP^+^ cells in control mice (PD-1 o.e.) and neutrophil depleted mice (PD-1 o.e. + dep.). **J**. FACS analysis of PD-1 overexpression in AT3 tumor cells. **K**. Average number of lung-associated GFP^+^ control and PD-1 overexpressing AT3 tumor cells in control and neutrophil depleted mice (n=5). *p<0.05, **p<0.01.

## DISCUSSION

In recent years our understanding of the multifaceted roles neutrophils play in the tumor microenvironment is growing, and they may either act to promote or limit tumor growth and progression. The present study identifies an important role of the PD-L1/PD-1 axis in regulating the direct anti-tumor function of neutrophils. The PD-L1/PD-1 axis is extensively explored in the context of cancer where tumor cell PD-L1 interacts with T cell PD-1 to limit adaptive anti tumor immune responses (**Fig. 5A**). However, in line with previous studies (He et al., 2015; Wang et al., 2017), we identified a PD-L1^pos^ subpopulation of neutrophils that is propagated in breast cancer. Thus far, the role of neutrophil PD-L1 was mainly reported in regard to the suppression of cytotoxic T cells (**Fig. 5B** (He et al., 2015)). Moreover, PD-L1^pos^ neutrophils were shown to contribute to the growth and progression in human tumors and are considered to be a poor prognostic factor indicative of reduced patient survival (Wang et al., 2017). In line with these studies, we observed a high expression of PD-L1 on the non-cytotoxic tumor infiltrating neutrophils (**Fig. 1C**). In contrast, cytotoxic neutrophils isolated from the circulation or the pre-metastatic niche are relatively low in PD-L1 expression (**Fig. 1B and 1D**). PD-L1^pos^ neutrophils exhibit a pro-tumor phenotype and can be mainly classified as immature LDN (**Fig. 1G**). This observation provides further support and possible mechanistic insight into the immunosuppressive LDN phenotype (Sagiv et al., 2015). Taken together, these observations suggest that PD-L1^pos^ neutrophils are characterized by having an immature, low-density, pro-tumor phenotype.

**Figure 5.**
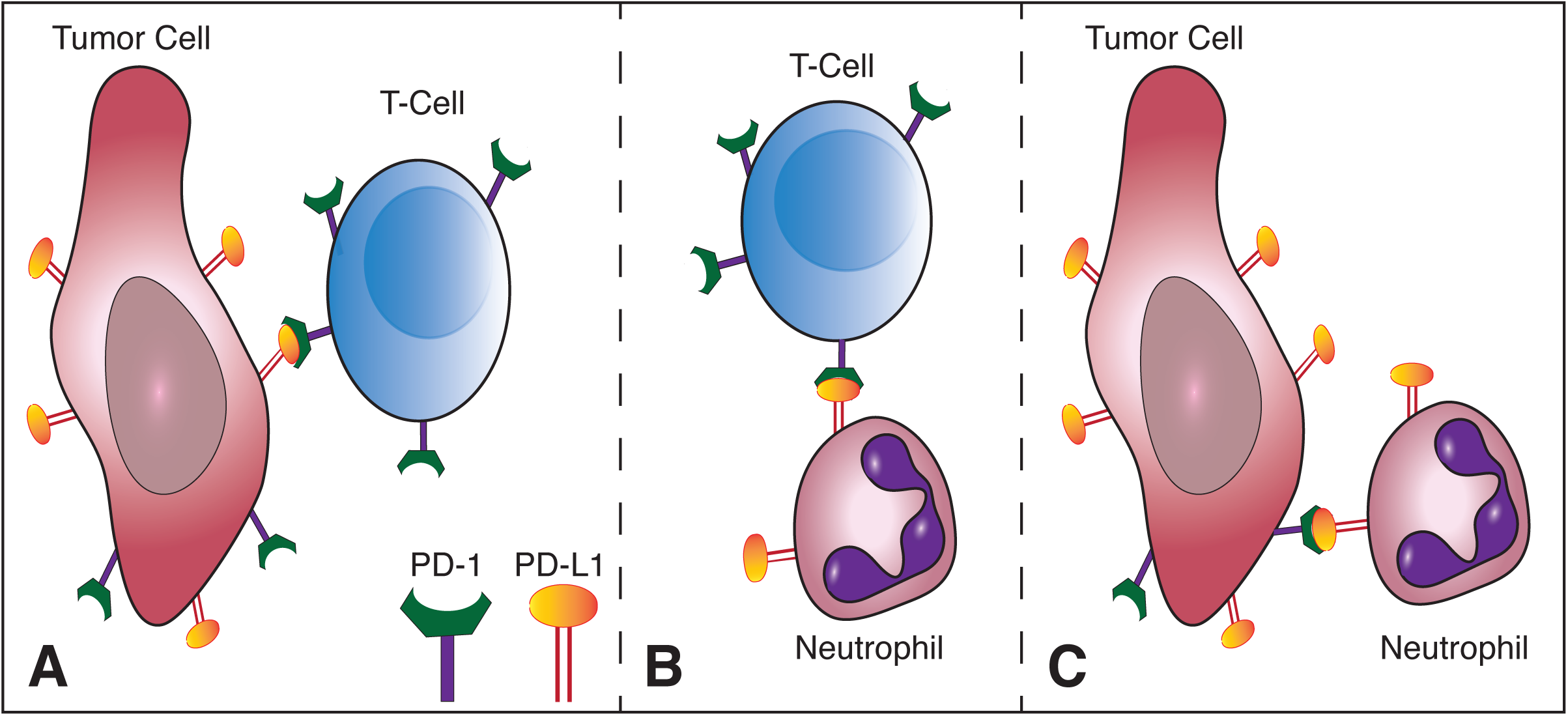
Mechanisms of tumor immune evasion mediated by the PD-1/PD-L1 axis. **A**. Tumor cell expressed PD-L1 serves as a ligand for PD-1 expressed on cytotoxic T Cells. This interaction leads to blocking of anti tumor T cell responses. Targeting this interaction serves as the basis for anti cancer immunotherapy. **B**. Neutrophils possessing immunosuppressive properties express PD-L1 which serves as a ligand for PD-1 expressed on cytotoxic T cells, blocking T cell anti tumor responses. **C**. Tumor cell expressed PD-1 interacts with PD-L1 expressed by neutrophils blocking cytotoxic anti tumor neutrophil responses.

The role of PD-L1 expressing neutrophils in the context of T cell suppression and tumor growth was already explored (**Fig. 5B**). However, the role played by the PD-L1/PD-1 axis in regulating anti-tumor properties of neutrophils was unknown. We and others (Kleffel et al., 2015) have shown that a subpopulation of tumor cells express PD-1 which may facilitate interactions with PD-L1 expressed by other cells. Accordingly we hypothesized that tumor cell PD-1 may interact with neutrophil PD-L1 and thus modulate neutrophil function. Indeed, blocking of the PD-L1/PD-1 axis in a co-culture setting enhanced tumor cell susceptibility to neutrophil cytotoxicity suggesting that the PD-1/PD-L1 axis is involved in regulating the direct neutrophil/tumor cell crosstalk. Moreover, this observation indicates that the PD-1/PD-L1 interaction in this context reduces the extent of neutrophil cytotoxicity (**Fig. 5C**). Since PD-1 expression levels vary among different tumors, we speculated that the effect different tumors have on the function of PD-L1^pos^ may correlate with the extent of PD-1 expression. To test this idea we overexpressed PD-1 in tumor cells and assessed tumor growth and progression mouse models of breast cancer. This led to a more aggressive phenotype as the tumors grew faster and had a higher metastatic load. Furthermore, neutrophil depletion dramatically enhanced metastatic seeding of control cells whereas PD-1 over-expressing tumor cells were not affected. Together, the observations implicate neutrophils in this process and suggest that PD-1 overexpressing cells are protected from neutrophils. Importantly, these experiments were done using immunodeficient NOD-SCID mice to eliminate the contribution of lymphocytes.

In summary, this study provides novel insight into the interaction of neutrophil PD-L1 with tumor cell PD-1 and the role it plays in regulating neutrophil cytotoxicity. We show that the PD-L1/PD-1 axis is also critical for the direct cytotoxicity, emphasizing the beneficial effect of blocking the PD-L1/PD-1 axis in cancer not only to enhance T cells functions, but also to enhance neutrophil anti-tumor activity.

## MATERIALS AND METHODS

### Animals

5-6-week-old BALB/c and NOD-SCID mice were purchased from the Harlan (Israel). All experiments involving animals were approved by the Hebrew University’s Institutional Animal Care and Use Committee (IACUC).

### Cell Lines

4T1 and AT3 mouse breast cancer cells were purchased from the ATCC and cultured in DMEM containing 10% heat-inactivated fetal calf serum (FCS). 4T1 and AT3 cells were transduced with a lentiviral vector (pLVX-Luc, MigR1-Luc or the triple-modality reporter gene TGL (Minn et al., 2005)) to stably express firefly luciferase. PD1kd 4T1 and AT3 cells were generated by lentiviral transduction with PD1 specific shRNAs from Sigma (TRCN0000097670-74). Control cells were transduced an empty vector (pLKO). For killing assay neutrophils and tumor cells were cultured in RPMI containing 2% heat-inactivated fetal calf serum (FCS).

### Plasmids

*Retroviral MigR1-Luc vector*: The open reading frame of Firefly luciferase was prepared from pLVX-luciferase using forward primer 5’-CAG TCC GCT CGA GGC CGC CAC CAT GGA AGA CGC CAA AAA CAT AAA G-3’ containing a XhoI restriction site, Kozac sequence followed by the ATG initiation code and reverse primer 5’-ACT TCC GGA ATT CTT ACA CGG CGA TCT TTC CGC CC-3’ containing an EcoRI restriction site and the stop codon and Phusion Flash High-Fidelity PCR master mix (Thermo Scientific). The amplified Luciferase PCR product was digested by XhoI and EcoRI and inserted into the XhoI/EcoRI site of the MigR1 plasmid.

#### Mouse PD1 expression vector

Mouse PD1 was prepared by PCR on cDNA from 4T1 breast cancer cells using Phusion Flash High-Fidelity PCR master mix (Thermo Scientific) and the following primer pair: F’ ACT TCC GGA ATT CGC CGC CAC CAT GTG GGT CCG GCA GGT ACC CTG GTC containing a EcoRI site and nt 67-89 of Pdcd1 (NM_008798.2) and R’ AAGGAAAAAAGCGGCCGCTCAAAGAGGCCAAGAACAATGTCC containing a NotI site and nt 907-930 of Pdcd1. The EcoRI/NotI-cut PCR product was mixed with annealed Flagx3 primers having sticky ends of EcoRI and NotI: F: ACT TCC GGA ATT CGC CGC CAC CAT GTG GGT CCG GCA GGT ACC CTG GTC and R’ AAG GAA AAA AGC GGC CGC TCA AAG AGG CCA AGA ACA ATG TCC; and inserted into the EcoRI/NotI site of the pLVX-Puro vector. The plasmid was sequenced at the Center for Genomic Technologies at the Givat Ram Campus, The Hebrew University of Jerusalem.

### Neutrophil Purification

Neutrophils were purified from orthotopically injected (mammary fat pad) 3-week 4T1 tumor-bearing mice as described before (Sionov et al., 2015a). In brief, whole blood was collected by cardiac puncture using heparinized (Sigma) syringe. The blood was diluted with 5 volume of PBS containing 0.5% BSA and subjected to a discontinuous Histopaque (Sigma) gradient (1.077 and 1.119). NDN were collected from the 1.077-1.119 interface. LDN were collected from the plasma-1.077 interface. Red blood cells (RBCs) were eliminated by hypotonic lysis. Neutrophil purity and viability were determined visually and were consistently >98% for NDN.

### Isolation of low and normal density neutrophils

Circulating NDN and LDN were purified from 4T1 tumor bearing mice using Histopaque gradient.

### Primary tumor and metastasis formation *in vivo* assay

NOD-SCID mice were injected into the mammary fat pad with 1×10^6^ control or PD1 over-expressing 4T1 cells. Tumors were measured weekly. On day 27 post tumor engraftment the mice were sacrificed and the lungs were excised for analysis metastatic spread. The lungs were sectioned with 100 µm intervals and stained with H&E. The number of metastatic foci was determined by histological examination. Data represents the average tumor size of at least 5 mice per group.

### Metastatic seeding assay

#### 4T1 metastatic seeding assay

Female NOD-SCID mice were orthotopically (mammary fat pad) injected with parental 4T1 tumor cells. On day 6 the mice were randomized and injected daily with either control IgG or neutrophil depleting anti-Ly6G (50 μ g/mouse i.p.). On day 8 the mice were injected to the tail vein with 5×10^5^ GFP^+^ cells. On day 10 the lung associated GFP^+^ cells were analyzed by fluorescent microscopy.

#### AT3 metastatic seeding assay

Female NOD-SCID mice were orthotopically (mammary fat pad) injected with parental 4T1 tumor cells. On day 14 the mice were randomized and injected daily with either control IgG or neutrophil depleting anti-Ly6G (50 μ g/mouse i.p.). On day 16 the mice were injected to the tail vein with 5×10^5^ GFP^+^ cells. On day 18 the lung associated GFP^+^ cells were analyzed by fluorescent microscopy.

### qPCR

Total RNA was isolated with TRI-Reagent (Sigma) according to the manufacturer’s instructions. RNA was reverse transcribed into cDNA using the AB high capacity cDNA kit (Applied Biosystems). 2 ng of converted cDNA was used for each reaction, using Kapa Sybr-Green Master Mix (Kapa Biosystems) and 300 nM of forward/reverse primer set. Analyses were done using the CFX384 BioRad Real Time PCR. GAPDH was used as a reference gene. The following primer sequences were used for gene expression analyses: GAPDH-F 5’-GCC TTC CGT GTT CCT ACC-3’, GAPDH-R 5’-CCT GCT TCA CCA CCT TCT T-3’,

### Peritonitis

BALB/c mice were injected i.p. with 1mg of Zymosan A from *Saccharomyces cerevisiae* (Sigma) diluted in 1ml of sterile PBS. Cells were retrieved using peritoneal lavage at four different time points (3, 24, 48, and 72 hours post i.p. injection). After purification and red blood cells (RBCs) hypotonic lysis the cell pellet was stained for PD-L1 and Ly6G and measured by flow cytometry.

### ROS assay

Purified PD-L1^pos^ and PD-L1^neg^ neutrophils were plated (2×10^5^) in 180 µl of HBSS (Biological Industries) in white 96-flat bottom wells (Corning) containing 50µM Luminol (Sigma). ROS production (chemiluminescence) and was measured using Tecan F200 microplate luminescence reader.

### *In Vitro* Killing Assay

Luciferase-labeled tumor cells (1×10^4^/well) were plated in 100 μl RPMI-1640 with 2% FCS in white 96-flat bottom wells (Corning). After 24 hours incubation, purified neutrophils (1×10^5^/well in 100 μl) were added to the plated tumor cells and co-cultured overnight in the presence or absence of anti-PD-L1 2μg (BioLegend). Following overnight incubation, luciferase activity was measured using Tecan F200 microplate luminescence reader. Extent of killing was determined by the ratio between tumor cells cultured alone and tumor cells co-cultured with neutrophils. *In vitro* experiments were repeated at least three times.

### PD-L1 positive and negative neutrophil isolation

PD-L1 positive and negative neutrophils were isolated using EasySep^TM^ PE positive selection kit (STEMCELL) according to the manufacturer’s instructions.

### BrdU labeling

Tumor bearing mice were injected i.p. with 100µL of BrdU solution (10mg/mL in sterile PBS, BD Pharmingen). Incorporation of BrdU was detected using the FITC BrdU Flow Kit (BD Pharmingen) and analyzed by flow cytometry.

### Antibodies

Mouse α-PD-L1 (BioLegend), α-mouse PD-L1-PE (BioLegend), α-mouse PD1-APC (BioLegend), α-mouse Ly6G-VioBlue (TONBO Biosciences), α-mouse Ly6G (clone 1A8, BioXCell).

### Statistical Analysis

For studies comparing differences between two groups, we used unpaired Student’s t tests. Differences were considered significant when p < 0.05. Data are presented as mean ± SEM.

